# The impact of ligand-induced oligomer dissociation on enzyme diffusion, directly observed at the single-molecule level

**DOI:** 10.1101/2024.01.28.577687

**Authors:** Yulia D. Yancheva, Saniye G. Kaya, Alexander Belyy, Marco W. Fraaije, Katarzyna M. Tych

**Author notes:** **Author Contributions**. These authors contributed equally (Yulia D. Yancheva, Saniye G. Kaya).

## Abstract

The existence of the phenomenon of enhanced enzyme diffusion (EED) has been a topic of debate in recent literature. One proposed mechanism to explain the origin of EED is oligomeric enzyme dissociation. We use mass photometry (MP), a label-free single-molecule technique, to investigate the dependence of the oligomeric states of several enzymes on their ligands. The studied enzymes of interest are catalase, aldolase, alkaline phosphatase and vanillyl-alcohol oxidase (VAO). We compared the ratios of oligomeric states in the presence and absence of substrate as well as different substrate and inhibitor concentrations. Catalase and aldolase were found to dissociate into smaller oligomers in the presence of their substrates, independently of inhibition, while for alkaline phosphatase and VAO, different behaviors were observed. Thus, we have identified a possible mechanism which explains the previously observed diffusion enhancement *in vitro*. This enhancement may occur due to the dissociation of oligomers through ligand binding.

---

The idea that some enzymes diffuse faster than expected from Brownian motion in the presence of their substrates has been a point of content in recent literature. It has been observed in some studies ^1–3^ and denied by others ^4,5^. Most experiments in which this phenomenon, termed enhanced enzyme diffusion (EED), has been observed, were conducted using fluorescence correlation spectroscopy (FCS) – an ensemble averaging technique requiring fluorescent labelling. However, Günther, et al. ^6^ discuss an artefact that arose in earlier reports of EED measured using FCS: the substrate of alkaline phosphatase, *p*-nitrophenyl phosphate (pNPP), was found to interact with the fluorescent dye, which was wrongly interpreted as EED. It is important to note that this artefact has only been observed in the case of alkaline phosphatase and its substrate pNPP, leaving the possibility of the existence of this effect in other enzymes open. The authors suggested that other single-molecule methods should be employed in order to further study the phenomenon.

EED may be biochemically relevant on various scales. While its existence poses fundamental questions relating to the structure and dynamics of individual enzymes, it may also be significant on the cellular level, in relation to recently described observations of the upper limit on Gibbs free energy dissipation in cells ^7,8^. This highlights its relevance on cellular level. However, its origin and mechanism are still unclear ^9^. Current EED models can be categorized as being catalysis-independent, such as the conformational changes an enzyme undergoes during ligand binding ^10,11^ and those where catalysis is essential for diffusion enhancement ^1–3^. One possible explanation for EED is based on a study by Jee, et al. ^12^ which suggests that oligomeric enzymes dissociate into their subunits in the presence of their substrate during catalysis. According to the Stokes-Einstein equation, the diffusion coefficient of a spherical particle is inversely proportional to its radius and therefore the subunits of an enzyme that has dissociated into smaller oligomers will diffuse faster than the enzyme in its larger, associated substrate-free state. Here, this mechanism is explored using mass photometry (MP). MP is a single molecule interferometric light scattering technique which can be used to determine the heterogeneity of a sample by measuring the masses of species present ^13^ (Figure 1). This technique can therefore be used to measure different oligomeric states of proteins. An advantage of MP is that measurements are performed in solution without fluorescent labelling. Furthermore, the concentrations used are in the nanomolar range, similar to those used in FCS. This makes MP a suitable technique to investigate EED under conditions close to those used in previous FCS measurements, but on the single-molecule level and under native conditions.

**Figure 1.**
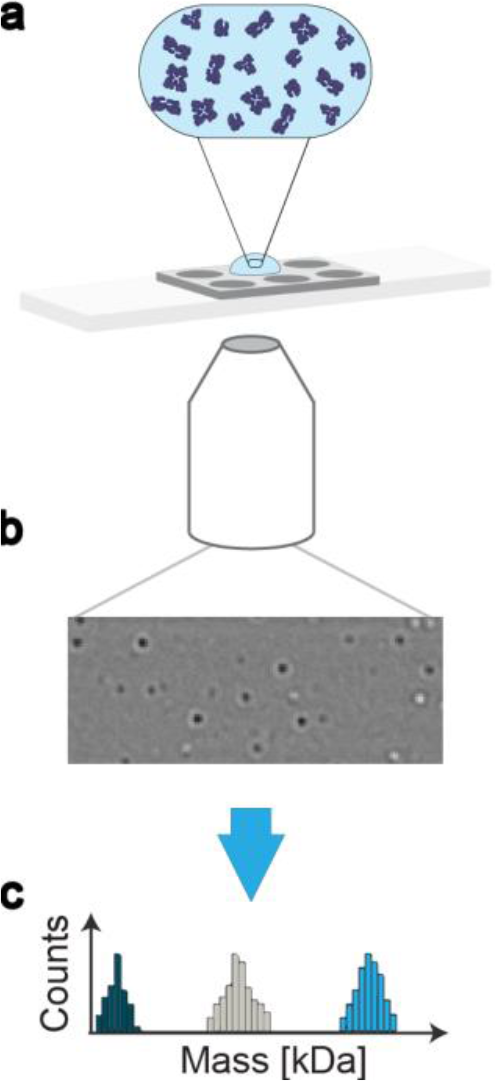
A mass photometry (MP) experiment. **a**. A glass slide with a gasket which contains six wells, each of which is used for a single measurement. For each measurement, 20 μL of diluted sample was used. **b**. The laser light goes through an interferometric microscope and the contrast change of each individual particle is measured. **c**. The contrast change is directly proportional to the mass of a particle. Mass histograms of the populations present in the sample are obtained.

The first enzyme we studied using MP was catalase (Figure 2a). Catalase is a fast exothermic enzyme that breaks down H_2_O_2_ to H_2_O and O_2_ and has been reported to exhibit diffusion enhancement up to 30 – 45% ^1,14,15^. The dissociation of oligomeric states of 50 nM catalase was investigated with between 0 and 50 mM of substrate. Upon addition of substrate, the enzyme dissociates into its subunits with a sharp change in states between 0 and 5 mM of substrate. The distribution of states (monomer, dimer, trimer, tetramer) remains constant from 5 to 50 mM H_2_O_2_, suggesting that the enzyme reaches an equilibrium state in terms of dissociation (Fig. 2b).

**Figure 2.**
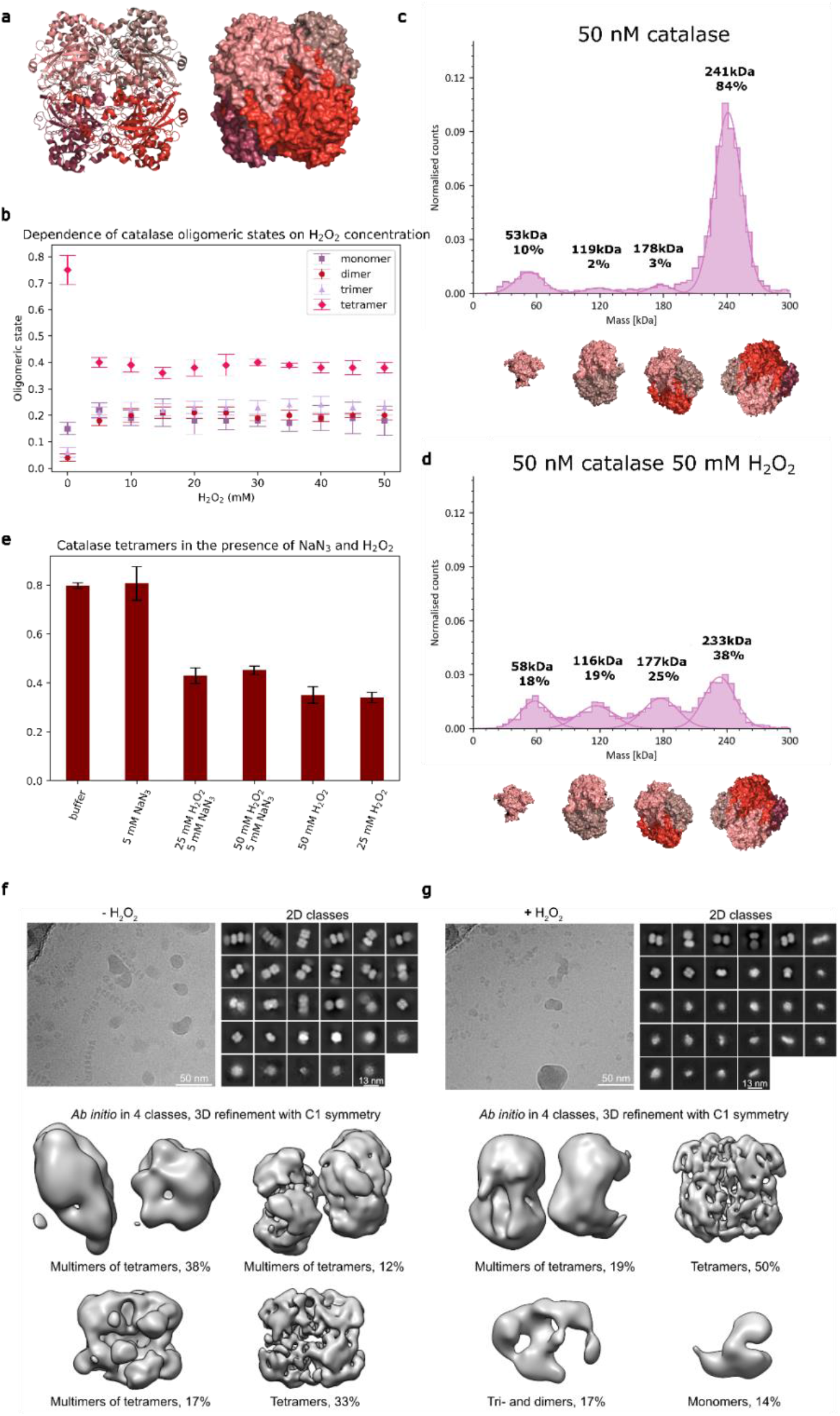
Oligomeric states of catalase (*Bos taurus*) in the presence of H_2_O_2_ (substrate) and NaN_3_ (inhibitor). **a**. Crystal structure of the apo state (PDB 1TGU_30_) in cartoon representation (left) and surface representation (right) of catalase. The different shades of red represent the monomers. **b**. Oligomeric states of catalase as a function of substrate concentration, where normalized oligomeric states as percentage of all oligomeric states are shown. **c**. Histogram of catalase in potassium phosphate buffer (pH 7.5) and **d**. in the presence of substrate. **e**. Catalase tetramers in the presence and absence of substrate and inhibitor. The percentage of tetramer population is shown on the y-axis. **f, g**. Cryo-EM micrographs, 2D class averages and low-resolution reconstructions of 4 μM catalase in the absence (f) or presence (g) of 50 mM H_2_O_2_ (substrate).

To investigate the effect of catalysis on oligomeric state, measurements were performed in the presence of H_2_O_2_ and the non-competitive inhibitor sodium azide (NaN_3_). The non-competitive inhibitor binds to the enzyme in such a way that it reduces the enzyme’s activity without affecting the binding of the substrate. To ensure that the inhibitor alone does not influence the oligomerization of the enzyme, a control experiment was performed (first two bars in the bar chart in Fig. 2e). The data presented in the middle two bars of the bar chart are for catalase with inhibitor and substrate, using different substrate concentrations while keeping the inhibitor concentration constant, while the last two bars represent catalase with just substrate. It can be seen that the amount of tetramer decreases upon addition of substrate, regardless of the presence of inhibitor, suggesting that catalysis is not required for the dissociation of the enzyme. To further confirm inhibition at these enzyme-to-inhibitor ratios, the activity of catalase was investigated in a bulk assay under the same conditions used in the MP experiments (Figures S2 and S3). This was further confirmed by the observation of bubbles, caused by the formation of dioxygen, which occurs only under turnover conditions (Figure S4).

A study on the temporal effects and reversibility of the dissociation of catalase was also performed. To study the reversibility of the dissociation, time course measurements of the system during catalysis were employed. As can be seen in figure S5, the dissociation of catalase is reversible and the oligomeric states return to the states prior to catalysis once the substrate is consumed. However, if an inhibitor is present and catalysis does not take place, the enzyme does not re-associate for at least 180 minutes, as can be seen in figure S6. This leads us to conclude that as long as a concentration of substrate that would trigger dissociation is present in the solution, no re-association will occur.

To ensure that our observations hold true for higher catalase concentrations, at which MP can no longer be used, we performed cryo-EM measurements. The oligomeric states of the enzyme at 4 μM were studied in the presence and absence of substrate as shown in figure 2 e, f. To ensure that the subunits do not re-associate in the time between the addition of the substrate and the freezing of the grid, 50 mM of the inhibitor, sodium azide was added in both conditions. The addition of inhibitor in the apo conditions shows that the dissociation is independent of the sodium azide and is only present upon addition of substrate. As can be seen in figure 2e, under apo-conditions catalase is either present as tetramer or forms higher-order structures as multimers of tetramers, previously termed ribbons ^16^. In the presence of 50 mM substrate (figure 2f), the populations of tetramers and multimers of tetramers decrease and a monomeric population appears as well as di- and trimeric populations. This observation is in line with the mass photometry data where the dissociation of tetramers is observed, thus providing evidence for the proposed EED mechanism of catalase on the single-molecule level and under native conditions.

The effects observed for catalase were also studied in other enzymes, one of which rabbit muscle aldolase. Aldolase is a slow endothermic enzyme with a K_M_ value of 0.013 mM for fructose-1,6-bisphosphate (FBP) ^17^. The enzyme has been reported to exhibit EED ^10^. While before this study it was hypothesized that exothermicity is necessary for EED and that a high turnover rate is required ^1^, the reports of aldolase exhibiting EED did not fit that hypothesis. Illien *et al*. ^10^ additionally report that the diffusion coefficient of aldolase increases in the presence of its competitive inhibitor pyrophosphate (PPi). It is important to note that as pointed out in ^15^, reports of EED of aldolase have been controversial across studies ^18^. In a study by Zhang *et al*. ^4^ using dynamic light scattering, no diffusion enhancement was observed in the presence of the substrate, FBP.

To investigate the mechanisms behind the observed EED for aldolase, the same experiment was conducted as for catalase. As can be seen in Figure 3, the tetrameric aldolase dissociates in a similar manner to catalase upon the addition of substrate, FBP, but plateauing at around 1 mM. In ^10^ it is discussed that dissociation of enzymes was not the reason for the observed EED since the diffusion goes back to base values after all substrate has been consumed. We hypothesize that the two observations are not mutually exclusive as the dissociated subunits are able to reassociate over time, as observed for catalase. To study the role of catalysis on the dissociation of aldolase, the competitive inhibitor PPi was used. A competitive inhibitor binds at the active site of an enzyme thus preventing the substrate from binding. Interestingly, in the presence of a competitive inhibitor and absence of substrate, partial dissociation of the tetramers into dimers and monomers is triggered (Fig. 3e).

**Figure 3.**
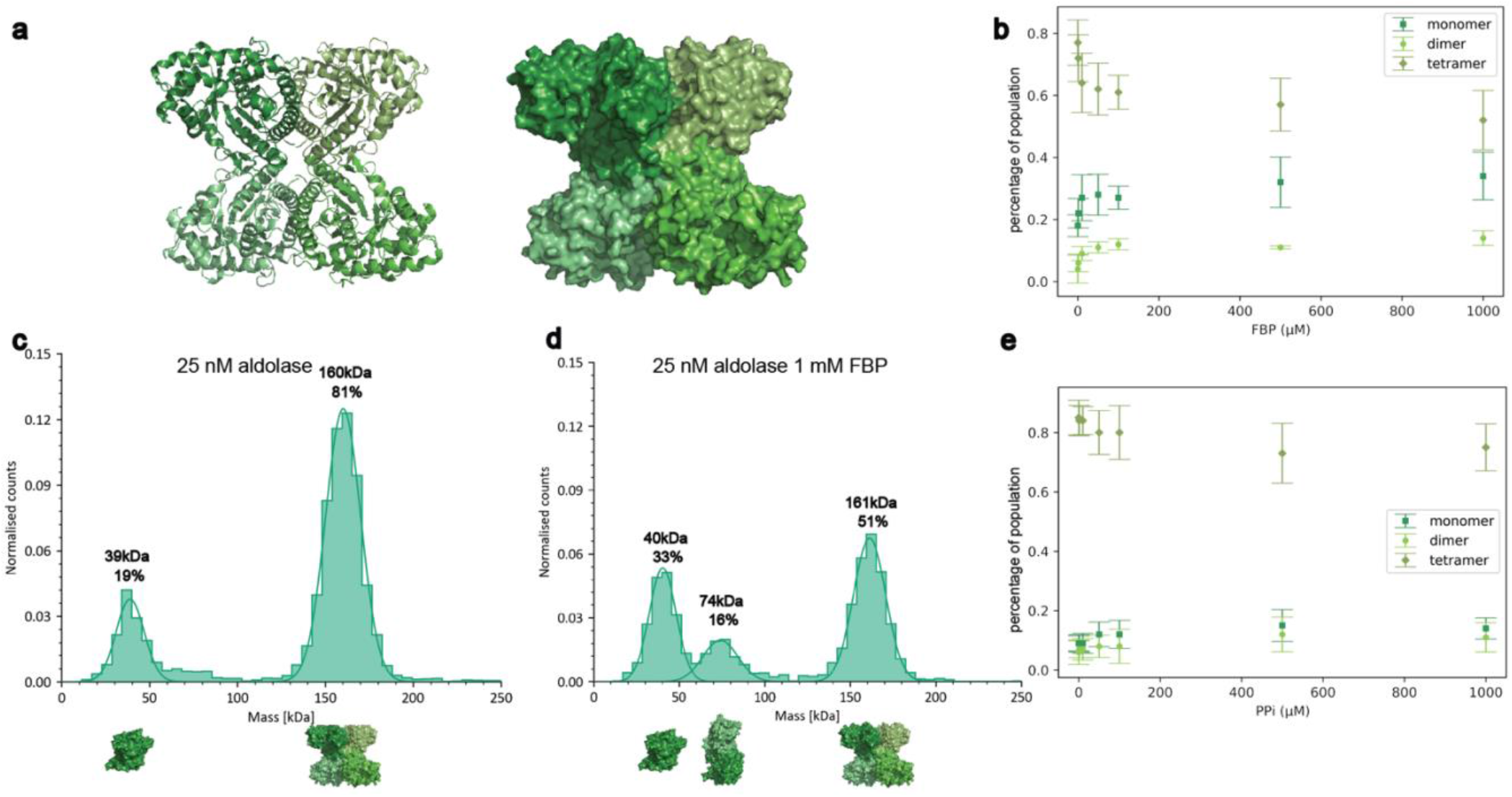
Oligomeric states of aldolase (*Oryctolagus cuniculus*) in the presence of FBP (substrate) and PPi (inhibitor). **a**. Crystal structure of the apo state (PDB 1ADO^31^) in cartoon representation (left) and surface representation (right). Each monomer is colored in a different shade of green. **b**. Normalized oligomeric states of aldolase as a function of substrate concentration, where normalized oligomeric states as percentage of all oligomeric states are shown. **c**. Histogram of aldolase in HEPES buffer (pH 7.4) and **d**. in the presence of substrate. **e**. Aldolase oligomeric states in the presence of inhibitor.

Another enzyme that was studied in the context of EED but sparked a lot of controversy in the field is alkaline phosphatase – a dimeric enzyme with a monomer mass of 57 kDa ^19^. In earlier research in enzyme diffusion, alkaline phosphatase (Fig. 4a) was reported to exhibit EED ^1^ but later papers show that what was interpreted as diffusion enhancement was an artefact related to the substrate interacting with the fluorescent labelling in FCS ^5^. While these findings were confirmed by Zhang et al. ^4^, the following measurements can be regarded as yet another confirmation of the artefact. In Fig. 4b it can be seen that there is no statistically significant change in the oligomeric states of 7.5 nM alkaline phosphatase when substrate is added, in the range 0 to 25 mM.

**Figure 4.**
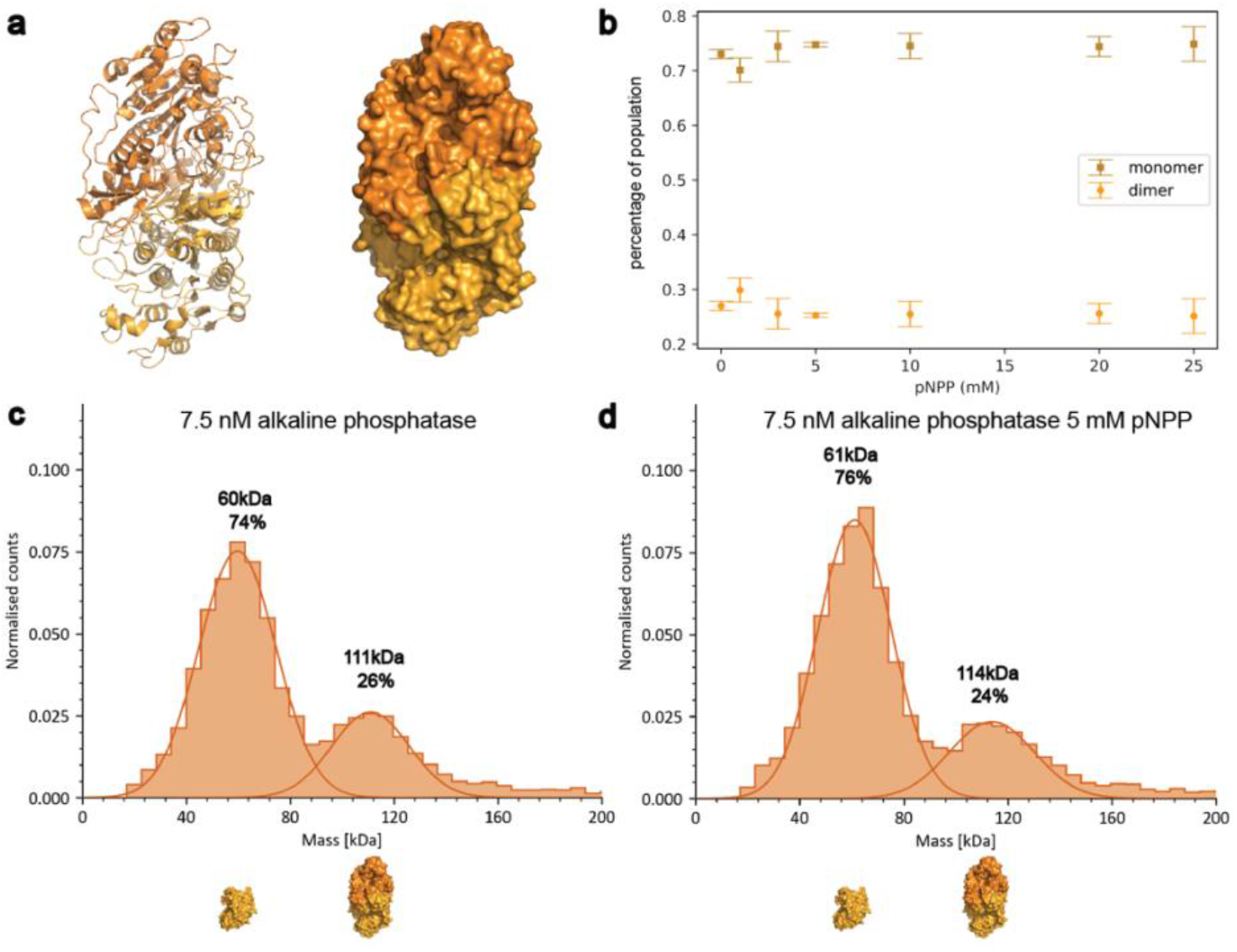
Oligomeric states of alkaline phosphatase (*Bos taurus*) in the presence of pNPP (substrate). **a**. Crystal structure of the apo state (PDB 1ALK_32_) in cartoon representation (left) and surface representation (right). The two shades of orange represent the two monomers that compose the dimer. **b**. Dependence of oligomeric states of alkaline phosphatase on substrate concentration. **c**. Histogram of alkaline phosphatase in alkaline buffer (see SI) and **d**. in the presence of 5 mM of substrate.

The fourth enzyme of interest in this study is VAO, which has not been studied in the context of EED. It is a fungal enzyme involved in the degradation of aromatic compounds and typically has an octameric structure consisting of eight identical subunits of 64 kDa (Fig. 5a), each harboring a covalently bound FAD cofactor ^20^. FAD binding stimulates the oligomerization of VAO into dimers and octamers, whereas the apo form of VAO consists of monomers and dimers ^21^. The most stable form of VAO is its octameric state, which can be regarded as a tetramer of dimers. It has been shown that also the dimers are active ^22,23^. These features make VAO a good candidate for studying dissociation and oligomerization in solution. As can be seen in figure 5b, VAO is mainly present as dimers and octamers. In the presence of substrate (vanillyl alcohol) the number of octamers increases at the expense of dimeric VAO, while the monomeric population remains constant (Figs. 5c and 5d). It has been observed before that the dimer-dimer interface is relatively small ^24^ and that changes such as the addition of chaotropic salt or cysteine oxidation can induce dissociation of octamers into dimers. Here, we show that interaction with substrate promotes association of dimers into octamers, rather than dissociation of higher oligomeric states to their subunits, as we observed for catalase and aldolase.

**Figure 5.**
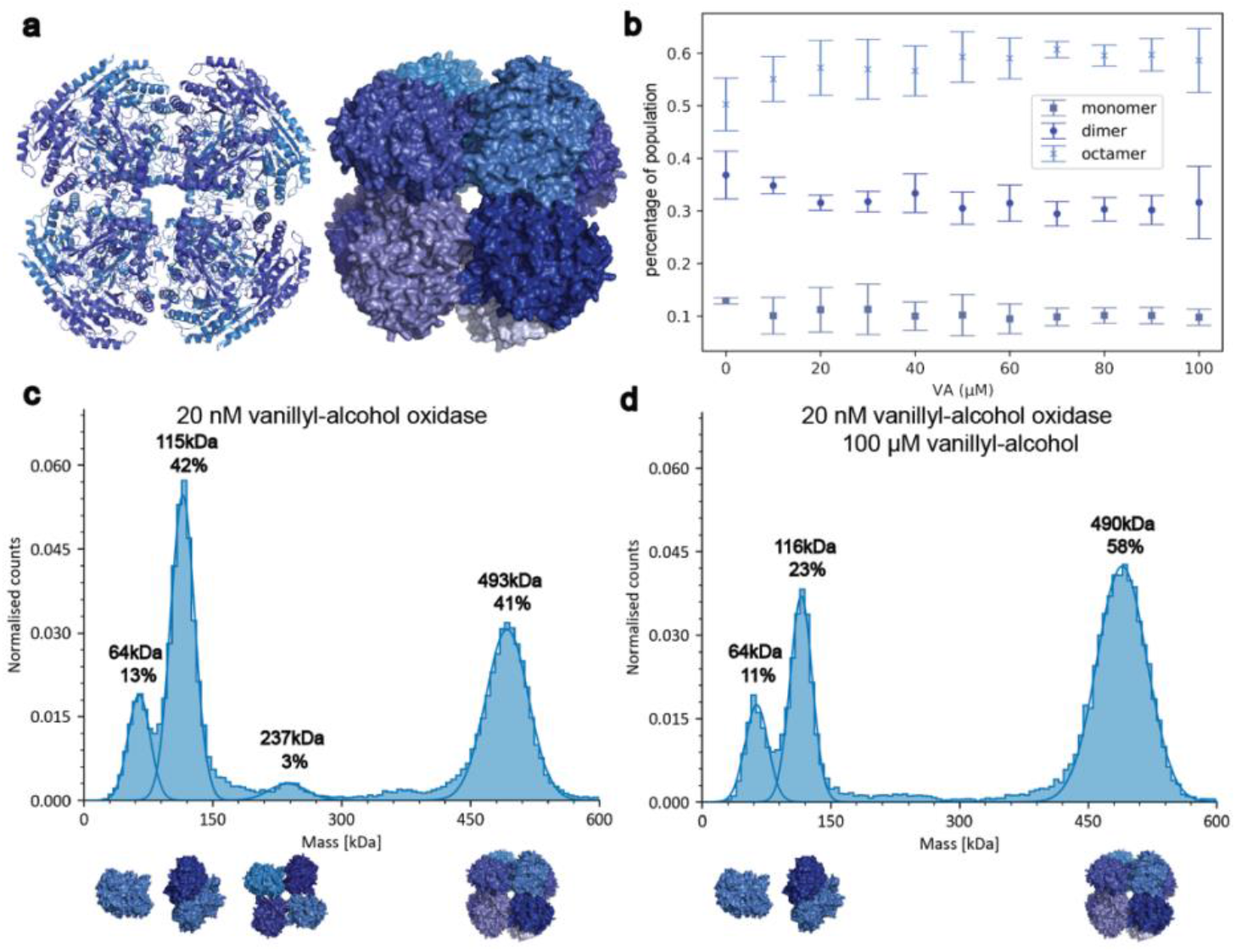
Oligomeric states of vanillyl-alcohol oxidase (VAO from *Penicillium simplicissimum*) in the presence of its substrate vanillyl-alcohol. **a**. Crystal structure of the apo state (PDB 1VAO_33_) in cartoon representation (left) and surface representation (right) The shades of blue represent individual monomers. **b**. Dependence of oligomeric states of VAO on its substrate concentration. **c**. Histogram of VAO in potassium phosphate buffer (pH 7.5) and **d**. in the presence of 100 μM vanillyl alcohol.

In this study we show that some of the enzymes that have previously been reported to exhibit EED, namely catalase and aldolase, dissociate in the presence of their substrates, while alkaline phosphatase does not dissociate in the presence of its substrate. Additionally, the oligomeric states of VAO were observed to shift towards an octameric state in the presence of its substrate, showing opposite behavior to that seen for catalase and aldolase. Catalase and aldolase were also studied in the presence of their respective inhibitors and it was observed that the dissociation of their tetramers does still occur, albeit to a lesser extent. It should be noted that since the measurements need to be performed in the nanomolar range for MP, the equilibrium ratios of the oligomeric states will differ compared to what would be observed at higher concentrations, for example in the micromolar range which is more relevant to biological processes ^25^, as shown by cryo-EM.

To further investigate the effects of the enzyme concentration on the dissociation and re-association of the proteins, simulations of this system at different concentrations should be performed. Moreover, molecular dynamics simulations and high-resolution cryo-EM should be employed to study the mechanisms by which ligands affect the quaternary structure of the enzymes and to provide additional insights into the changes of the monomer interfaces after dissociation.

While oligomerization is essential for protein function, structural factors driving this oligomerization remain unclear for many proteins ^26^. Previously it was reported that substrate binding influences oligomerization of an enzyme ^27^ and that oligomerization can play a crucial role in enzymatic activity ^28^, yet we observe oligomer dissociation of catalase and aldolase upon ligand binding. This observation poses questions regarding the oligomerization in the first place and the need for its reversibility. We hypothesize that this dissociation may have a regulatory role. Another possible benefit for enzyme dissociation was discussed by Agudo-Canalejo et al.^29^, suggesting that enzyme dissociation can be beneficial for proteins to find their target and react more rapidly.

We conclude that enzyme dissociation is one likely mechanism for previously reported EED. The dissociation is not catalysis driven and should be studied in context of the changes of the enzymes’ structures and their oligomerization interfaces. Thus, ligand binding may play a dominant role in the hydrodynamic properties of enzymes, while catalysis may be less relevant for EED.

## ASSOCIATED CONTENT

### Supporting Information

The Supporting Information is available free of charge on the ACS Publications website: Preparation and characterization of enzymes, mass photometry, additional data for hexokinase and aldolase, diffusion enhancement calculations.

## AUTHOR INFORMATION

## Notes

The authors declare no competing financial interests.

## ACKNOWLEDGMENT

YDY and SGK are supported by the NWO NWA grant number NWA.1292.19.170, “Limits to Growth”. We thank Ronald van der Meulen for insightful discussions.

## SYNOPSIS TOC

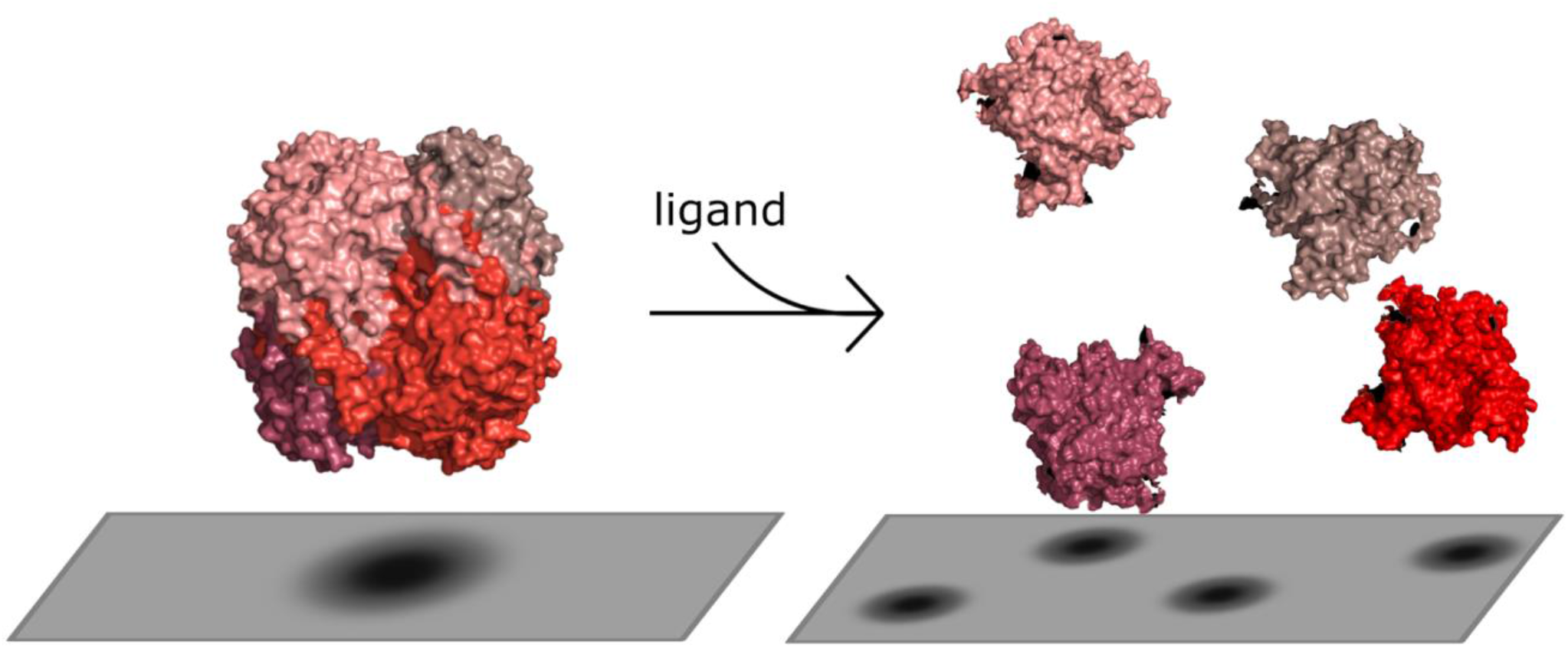

