## Supplemental Data and Figures for "The impact of ligand-induced oligomer dissociation on enzyme diffusion, directly observed at the single-molecule level"

### Supplementary information

#### 1 Preparation and characterisation of enzymes

There were four enzymes investigated in this paper. The first enzyme studied is catalase from bovine liver, a tetrameric enzyme with $K_m$ = 28 mM, that breaks down $H_2O_2$ to $H_2O$ and $O_2$ [1] [2]. It is important to note that $K_m$ in the case of catalase is not the real $K_m$ because the active site of catalase is not saturable via $H_2O_2$ [3]. The second enzyme studied is aldolase from rabbit muscle, another tetrameric enzyme with $K_m$ = 0.013 mM that converts fructose-1,6-bisphosphate (FBP) into D-glyceraldehyde 3-phosphate (G3P) and dihydroxyacetone phosphate (DHAP). Alkaline phosphatase from bovine intestinal mucosa was also studied, it is a dimeric enzyme with $K_m$ = 0.62 mM that converts pNPP to p-nitrophenol (PNP). The fourth enzyme studied is vanillyl-alcohol oxidase (VAO) from *P. simplicissimum*, an octameric enzyme [4] with $K_m$ = 0.04 mM that oxidises vanillyl alcohol (VA) to vanillin. Catalase [5], aldolase [6] and alkaline phosphatase [1] have been reported to exhibit EED, while VAO has not been studied in this context. However, since it is an octameric enzyme this makes it a control to confirm the reliability of the experimental method. While the diffusion enhancement of alkaline phosphatase has been attributed to an experimental artefact, we deemed important to include it in this study as an additional control to confirm the findings of latest research on this enzyme.

Catalase from bovine liver (9001-05-2), fructose bisphosphate aldolase from rabbit muscle (9024-52-6), alkaline phosphatase from bovine intestinal mucosa (9001-78-9) and Hexokinase I from *Saccharomyces cerevisiae* (9001-51-8) purchased from Sigma-Aldrich. Since the catalase is offered as suspension in water containing 0.1% (w/v) thymol, it had to be pelleted by centrifugation. To do so it was centrifuged at 13g for 5 min at 4°C. The thymol solution was taken out and the palette was dissolved in water and centrifuged again at 13g for 5 min at 4°C. The water was taken out and replaced with buffer. It was then gently mixed and placed in a tabletop heater at 30°C for 5 min, mixed again and put back in the tabletop heater. This was repeated until the pellet was fully dissolved. The buffer used for the washing of catalase is potassium phosphate buffer pH 7.5 and the same buffer was used for the VAO measurements. For aldolase, 50 mM HEPES pH 7.4 was used. The buffer in which alkaline phosphatase was diluted is alkaline buffer (20 mM TRIS, 100 mM NaCl, 1 mM MgCl, 20 µM $ZnCl_2$) pH 8.5.

##### 1.1 Expression and purification

Octameric vanillyl alcohol oxidase from *P. simplicissimum* was expressed and purified according to the previous reports [7] [8].

##### 1.2 Steady state kinetics

For each enzyme, steady state kinetics was performed in a substrate range matching with mass photometry experiments to see that the enzymes are active in the measurements. All steady state kinetics were performed by using V-660 spectrophotometer, Jasco. The activity of aldolase and alkaline phosphatase was measured by following previous studies [6] [5].

Catalase activity was measured by following the depletion of $H_2O_2$ at 240 nm. Reaction rate was calculated using extinction coefficient $\epsilon = 43.5m^{-1}cm^{-1}$ at 240 nm [5].

The rates were calculated by the slope of the curves observed. The data was fitted to the Michaelis-Menten equation $Y = \frac{k_{cat}*X}{K_M+X}$, where all symbols have their usual meanings, to determine the corresponding $K_m$ and $k_{cat}$ values.

#### 2 Mass photometry

All mass photometry measurements we performed using the Refeyn TwoMP mass photometry setup. All measurements were recorded using the AcquireMP software. Microscope coverslips sized at 24 mm x 50 mm (VMR) were cleaned by being immersed in 3% mucasol solution for at least 30 min. Subsequently, these coverslips underwent a thorough washing process involving sequential rinses in HPLC-grade ethanol and Milli-Q $H_2O$, repeated at least three times for each solvent. The coverslips were then meticulously dried using nitrogen flow.

To set the experimental stage, a 6-well silicon gasket was positioned onto a clean coverslip, which was then carefully inserted into the instrument. The protein samples underwent necessary dilution just before the measurement session, achieving a working concentration in the nanomolar range. In each measurement cycle there was a control measurement of enzyme in its corresponding buffer. To perform the measurement, $10\mu L$ of buffer, which had been filtered through a 0.22-$\mu$m syringe, was added into the designated well. In the case of measurements with substrate, $10\mu L$ of substrate was added instead. Then the focus was found using the droplet dilution setting of the AcquireMP software. Upon achieving focus, 10 $\mu L$ of the protein sample was introduced into the same well and gently mixed. Following mixing, data collection was initiated and continued for a duration of one minute. The same procedure as for substrate was followed with the competitive inhibitor, PPi, of aldolase. In the case of catalase, since the measurements were performed with non-competitive inhibitor, the enzyme was first incubated with the inhibitor for one minute and subsequently added to either buffer or substrate, depending on the type of measurement performed.

Subsequent data processing and analysis were conducted using the DiscoverMP software, also developed by Refeyn (UK).

#### 2.1 Mass calibration

Mass calibration was performed using protein standards of known masses. The same mass calibration was applied on the data presented in the main text. For aldolase and alkaline phosphatase the calibration obtained using the masses of monomeric and dimeric eugenol oxidase (*P. simplicissimum*, 59 kDa and 117 kDa, respectively) and tetrameric catalase (*Bos taurus*, 240 kDa) as standards. For VAO the mass calibration applied was obtained using the 66 kDa, 480 kDa and 1048 kDa peaks from NativeMark™ Unstained Protein Standard (ThermoFisher, catalog number LC0725). The calibration for catalase was obtained using the four peaks (60 kDa, 120 kDa, 180 kDa and 240 kDa) of OpOx. In this study we are not investigating the masses of the proteins but the ratios of their oligomeric states. Since the masses of all proteins used are already known and reported, a mass calibration was applied only for visual clarity.

#### 2.2 Data collection

Instead of looking at the absolute value of the molecules of each oligomeric state, the parameter investigated in these experiments was the normalised number of counts, presented as a percentage of the total events of interest. Normalisation was required to eliminate any artefacts arising from events which are not of interest, such as noise or higher order oligomeric structures. This way the oligomeric states are compared relative to each other and not relative to the total number of events detected.

The number of particles detected per oligomeric state varies stochastically throughout the measurements due to the short measurement time of 60 s. To account for this effect, each data point represents the mean of three measurement days and the error is represented by the standard deviation of their means. For each measurement day the enzymes and substrates are diluted from stock in the same dilution steps to additionally account for pipetting error which can have significant effect at the nanomolar concentration range in which measurements are performed. To further account for the variability of the events detected per measurement, a data point from each measurement day consists of the merging of at least three individual measurements at the same conditions. The repeats of the measurements per condition are taken within two to three hours. In order to exclude the possibility of dissociation occurring due to the enzymes being kept at low concentration for extended time periods before measurement, we ensured that the repeats of each condition were not performed consecutively. This way the conditions without ligand were spaced over time across the whole period in which the measurements were performed.

As mentioned above, catalase breaks down $H_2O_2$ to $H_2O$ and $O_2$. In this reaction, oxygen bubbles are released, which has raised concerns in previous research [9]. Although it has been discussed that bubble propulsion is not a possible explanation of the diffusion enhancement observed [10], it is essential to consider if these bubbles could interfere with the measurements. To address these concerns, a video of an oxygen bubble interfering with a measurement of catalase has been included.

Data from videos in which an oxygen bubble is observed have been discarded during the data analysis process.

### 3 Cryo-EM

Samples of 4 $\mu$M catalase mixed with 50 mM sodium azide in the presence of absence of 50 mM $H_2O_2$ were applied onto a freshly glow-discharged copper R1.2/1.3 300 mesh grid (Quantifoil), blotted for 3 s on both sides with blotting force 0 and plunge-frozen in the liquid ethane-propane mixture using the Vitrobot Mark IV system (Thermo Fisher Scientific) at 13 °C and 100% humidity. Datasets of 355 (with substrate) and 402 (without substrate) micrographs were collected using a Talos Arctica transmission electron microscope (Thermo Fisher Scientific) equipped with an XFEG at 200 kV using the automated data-collection software EPU version 2.7 (Thermo Fisher Scientific). 1 image per hole with defocus range of -2 - -3 $\mu$m were collected with K2 detector (Gatan) operated in counting mode. Image stacks with 50 frames were collected with the total exposure time of 9 sec and total dose of 50 $e^-/Å^2$. After motion correction and CTF estimation performed using MotionCorr2 [11] and CTFFIND 4.1 [12] in Relion version 3.1 [13], particles were picked using crYOLO [14] with the general picking model and low confidence threshold of 0.1 to ensure that all particles are picked. A stack of 21986 (with substrate) and 19146 (without substrate) particles was imported in CryoSPARC and subjected to 2D classification into 50 classes. Only clear non-protein particles were discarded and the remaining 13786 (with substrate) and 13781 (without substrate) particles were subjected to ab-initio reconstruction with simultaneous sorting in 4 classes, followed by non-uniform refinement.

### 4 Additional data

#### 4.1 Hexokinase

Hexokinase from *Saccharomyces cerevisiae* is also mentioned as one of the enzymes that exhibit EED [5]. Hexokinase is a dimeric enzyme with monomer mass of 55 kDa. In order to investigate the dissociation of hexokinase upon catalysis using mass photometry, it is important to work in the nanomolar range. However, since the dissociation constant of hexokinase is in that range, once the enzyme is diluted, there are only monomeric species present in the solution, as can be seen in Figure S1. This result has been discussed in previous studies: [5] also reported that although the enzyme is dimeric, in FCS experiments, the obtained diffusion coefficient corresponds to the one of a monomeric hexokinase. This is likely due to the high dissociation constant of the enzyme [10] [15] and the low concentrations that are used in FCS measurements, which are in the same range as the concentrations used in MP. Further measurements of the oligomerization dynamics of this enzyme could not be performed at this concentration.

#### 4.2 Aldolase

To comment on the role of catalysis in the dissociation, one should compare figures 3b and 3e. While the dissociation of aldolase in the presence of PPi has been observed, it is to a lesser extent than in the case with FBP. Additionally, the comparison between the two conditions should also be viewed in the context of the data analysis process - each point on these graphs is the mean of three individual measurements and the error represents the deviation of the values from the mean. While in both cases the error bars are larger than in the case of catalase (Figure 2b), this error does not only account for random error but also for systematic error. Since the same analysis method was employed for all four enzymes for consistency, it is important to consider the analysis of each individual measurement day. From 7, it can be seen that while the trend of aldolase dissociating in the presence of FBP is the same for all measurements, the absolute percentages can differ. We observe a similar dissimilarity between the data sets for aldolase in the presence of PPi: in some of the data sets the enzyme falls apart in a similar manner to the way it does in the presence of FBP while in other data sets, such dissociation is not observed. We noticed this behavior and ensured to always dilute the enzyme and substrate using the same dilution steps and while the repeatability was improved, the behavior in some of the data sets remains unexplained at this stage. If data for each measurement day is plotted separately (7), it can be seen that the dissociation still occurs but not as consistently as in the case of catalase. This leads us to hypothesize that the quaternary structure is weakened in the presence of inhibitors, but that weakening is not necessarily enough to lead to the dissociation of the enzyme. With this observation of the inconsistency of the behavior of aldolase, we suggest that the contradicting reports of EED [6] [16] [1] [17] for aldolase could be due to the lack of repeatability that we observe in our measurements.

### 5 Diffusion enhancement calculations

To estimate the average diffusion enhancement of that would be observed if catalase dissociates in the same ratios as observed in the mass photometry experiments and compare it we did the following calculations.

From Stokes-Einstein equation the diffusion coefficient can be calculated as a function of the hydrodynamic radius of a particle

$$D = \frac{k_b T}{6\pi\eta R} \tag{1}$$

The average experimentally measured diffusion coefficient $D_{exp}$ can be estimated by looking at the amount of each oligomeric state present in the solution.

$$D_{exp} = a\frac{k_b T}{6\pi\eta R_M} + b\frac{k_b T}{6\pi\eta R_D} + c\frac{k_b T}{6\pi\eta R_{Tr}} + d\frac{k_b T}{6\pi\eta R_T}, \tag{2}$$

where $R_M$, $R_D$, $R_{Tr}$ and $R_T$ are the radii of the monomer, dimer, trimer and tetramer respectively, assuming spherical particles. The coefficients $a, b, c$ and $d$ are the percentages of total population for monomer, dimer, trimer and tetramer respectively. Additionally, since all four subunits are equal in size, we can assume that

$$R_D = 2R_M, R_{Tr} = 3R_M \text{ and } R_T = 4R_M. \tag{3}$$

Equation 2 can be rewritten as

$$D_{exp} = a\frac{k_b T}{6\pi\eta R_M} + b\frac{k_b T}{6\pi\eta 2R_M} + c\frac{k_b T}{6\pi\eta 3R_M} + d\frac{k_b T}{6\pi\eta 4R_M} \tag{4}$$

or

$$D_{exp} = \frac{k_b T}{6\pi\eta R_M}(a + \frac{b}{2} + \frac{c}{3} + \frac{d}{4}) = \frac{1}{12}\frac{k_b T}{6\pi\eta R_M}(12a + 6b + 4c + 3d)) \tag{5}$$

Let the experimental diffusion coefficient in the absence of substrate be $D_0$ and the one in the presence of substrate be $D_1$. Since we are interested in the ratio of the diffusion coefficients rather than their absolute values, we can write

$$\frac{D_1}{D_0} = \frac{\frac{1}{12}\frac{k_b T}{6\pi\eta R_M}(12a_1 + 6b_1 + 4c_1 + 3d_1)}{\frac{1}{12}\frac{k_b T}{6\pi\eta R_M}(12a_0 + 6b_0 + 4c_0 + 3d_0)} = \frac{12a_1 + 6b_1 + 4c_1 + 3d_1}{12a_0 + 6b_0 + 4c_0 + 3d_0} \tag{6}$$

Taking the values of the coefficients for catalase from the MP experiments (Fig 2c, 2d) in percentage where $a_0 = 0.1, b_0 = 0.02, c_0 = 0.03, d_0 = 0.85$ and $a_1 = 0.18, b_1 = 0.19, c_1 = 0.25, d_1 = 0.38$.

$$\frac{D_1}{D_0} = 1.36 \tag{7}$$

which corresponds to the reports of 30-45% enhancement as discussed in the main text.

### 6 Figures

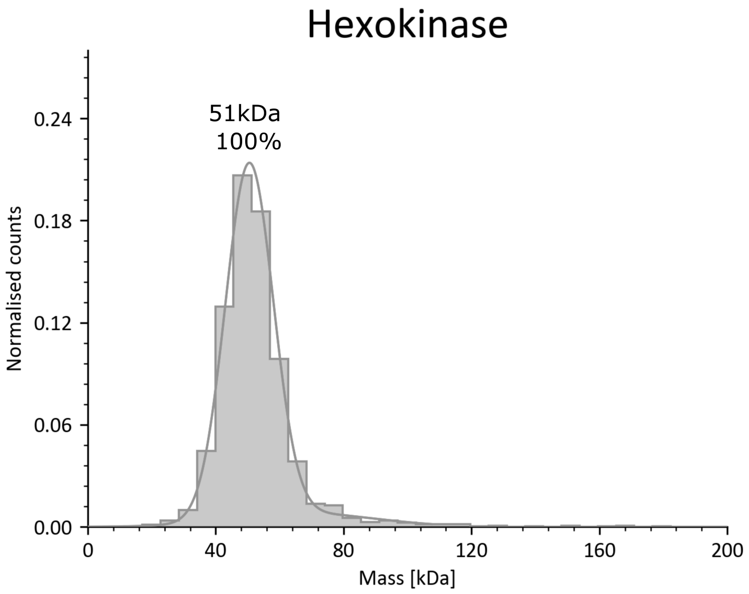


Figure 1: Mass photometry histogram of hexokinase from *Saccharomyces cerevisiae* in PBS buffer (pH 7.5). On the x-axis the mass is presented and on the y-axis are the normalized counts. Hexokinase is expected to be dimeric but due to its high dissociation constant in the concentration range accessible for the mass photometry it only exists in monomeric form.

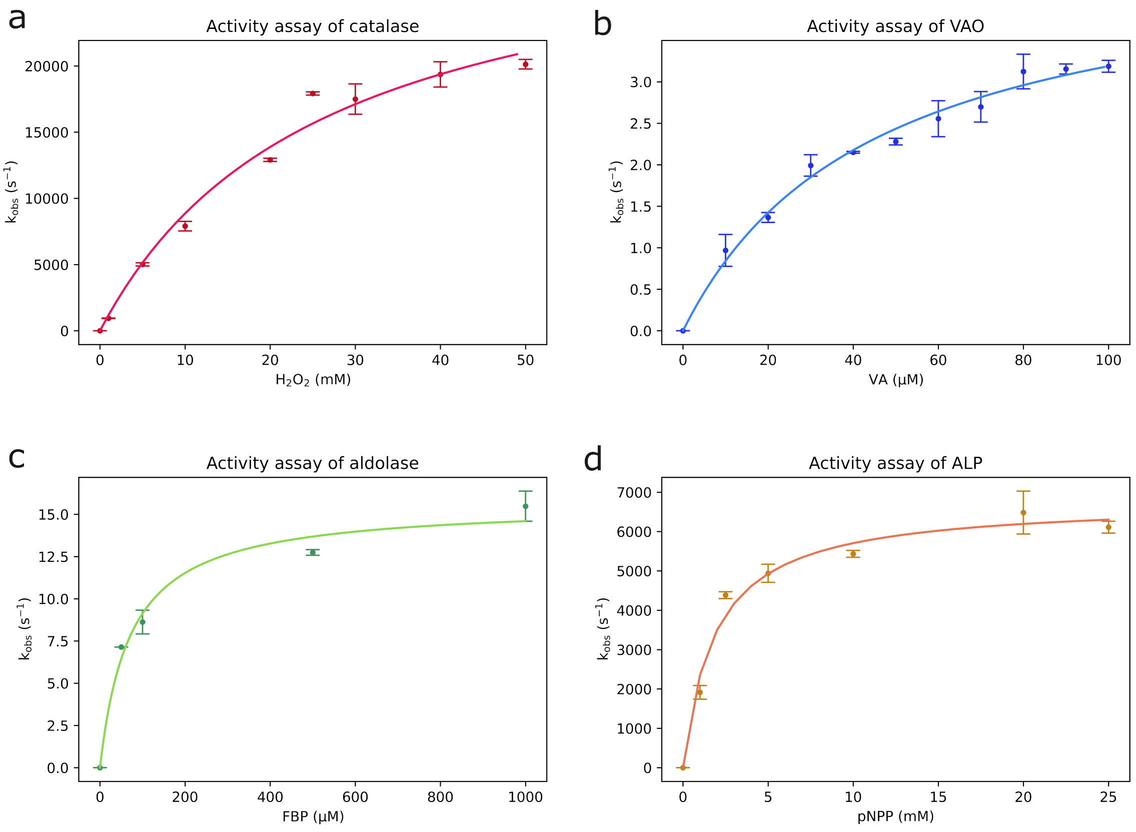


Figure 2: Michaelis-Menten fit of the values obtained from enzyme activity assay. a. Catalase b. vanillyl alcohol oxidase c. aldolase and d. alkaline phosphatase. Error bars represent the standard deviation of triplicates.

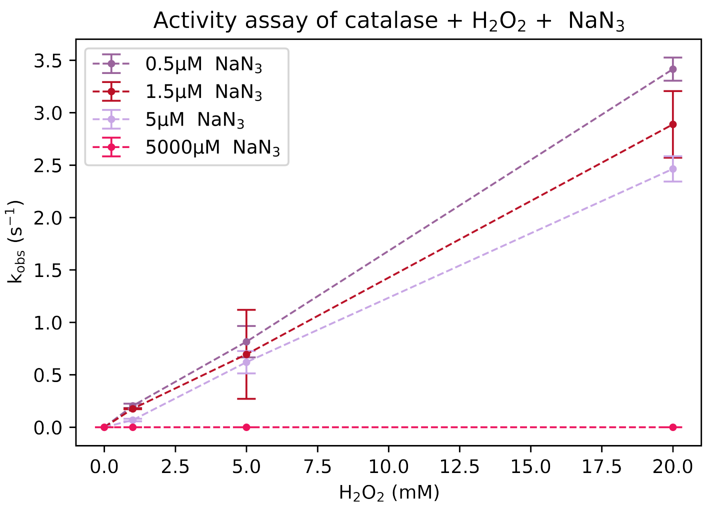


Figure 3: Activity assay of catalase in different $NaN_3$ concentrations. For 5 mM of $NaN_3$ no activity was observed. Dashed lines are for visual clarity.

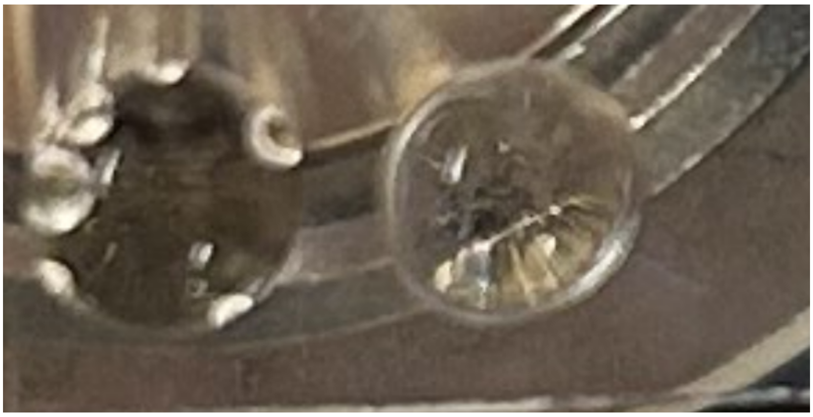

Figure 4: Two wells on a mass photometry gasket containing catalase in the presence of 50mM $H_2O_2$ in the absence (left) and presence (right) of $NaN_3$.

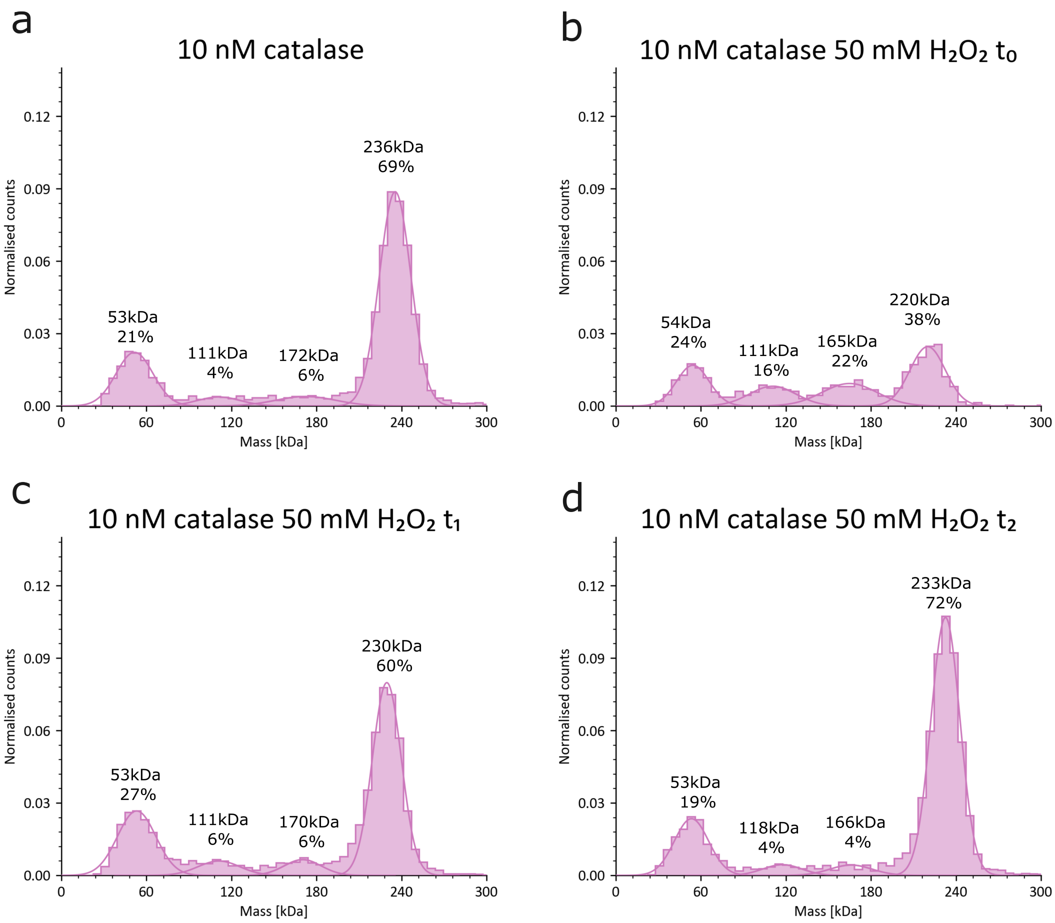


Figure 5: Reversibility of the dissociation of catalase. a. Catalase oligomeric states in apo conditions b. Catalase oligomeric states at $t_0 = 30 \pm 3$ s after addition of substrate c. Catalase oligomeric states at $t_1 = 143 \pm 1$ s d. Catalase oligomeric states at $t_2 = 265 \pm 10$ s

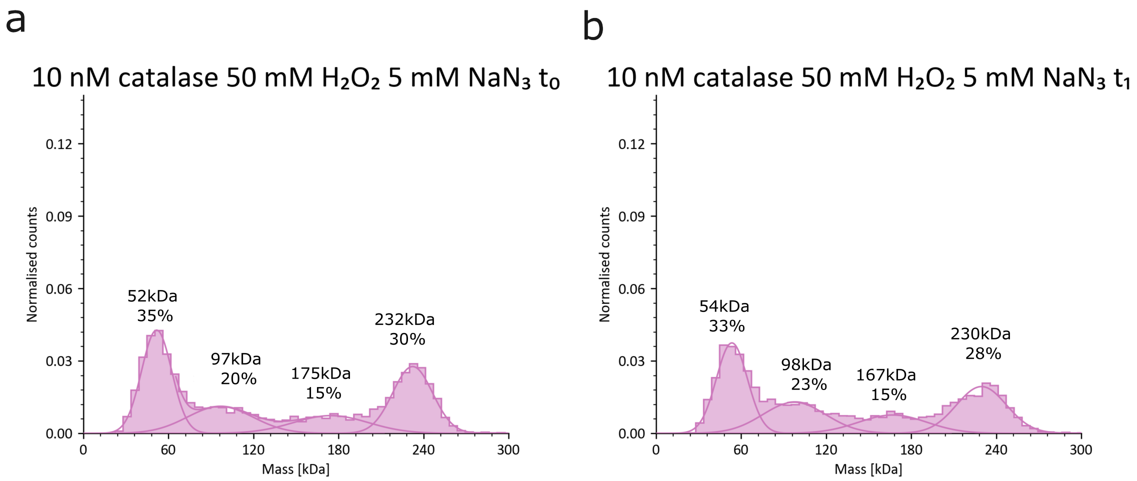


Figure 6: No reversibility of the dissociation of catalase in the presence of inhibitor. a. Catalase oligomeric states upon addition to substrate and inhibitor b. Catalase oligomeric states at $t_1 = 180$ min after addition of substrate and inhibitor

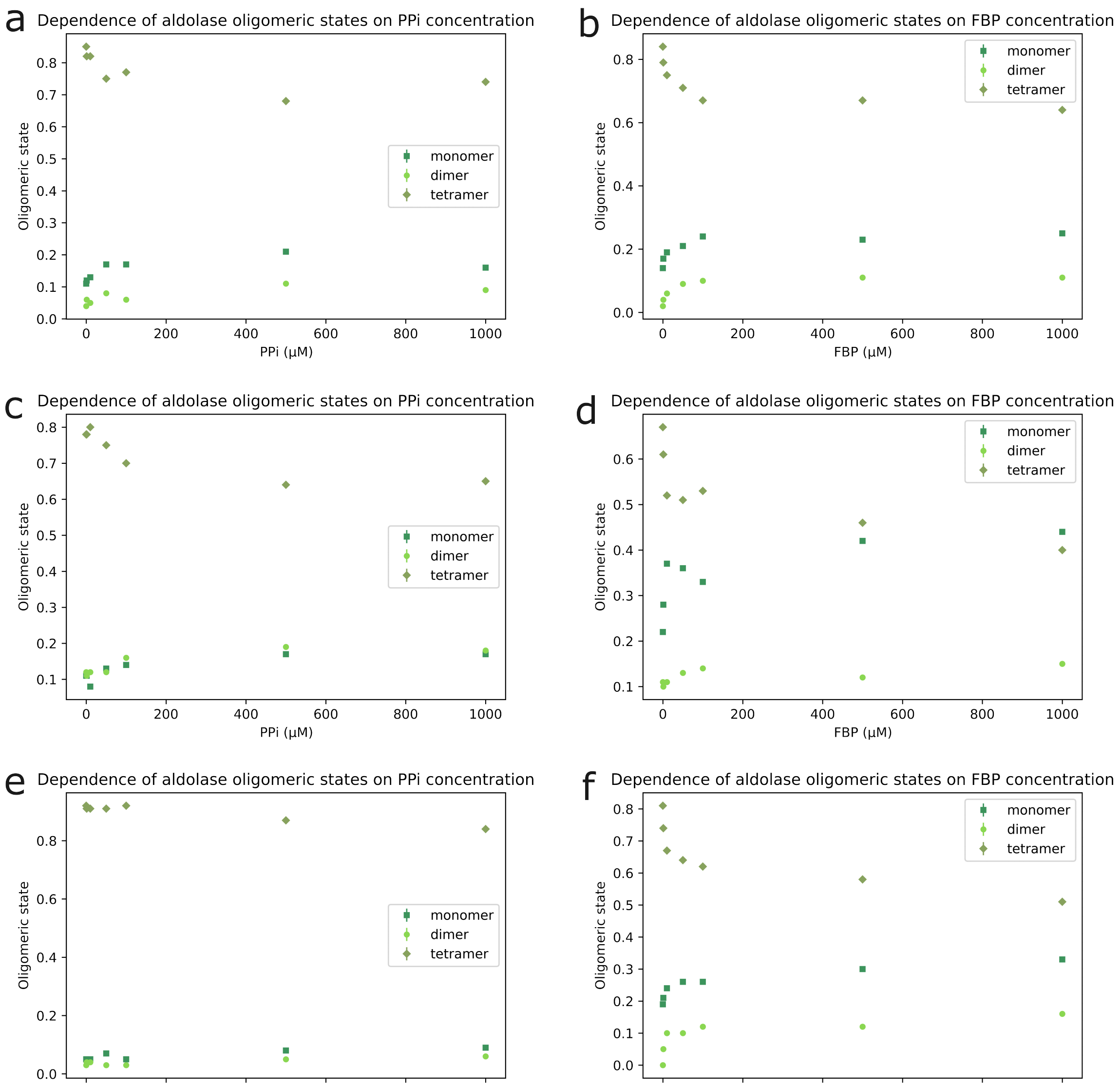


Figure 7: Individual measurements of the oligomeric states of aldolase in the presence of its inhibitor PPi (a, c, e) and in the presence of its substrate FBP (b, d, f)
